# Mechanism of an Intrinsic Oscillation in Rat Geniculate Interneurons

**DOI:** 10.1101/2024.06.06.597830

**Authors:** Erica Y. Griffith, Mohamed ElSayed, Salvador Dura-Bernal, Samuel A. Neymotin, Daniel J. Uhlrich, William W. Lytton, J. Julius Zhu

**Affiliations:** Department of Neural and Behavioral Sciences, SUNY Downstate Health Sciences University, Brooklyn, NY; Center for Biomedical Imaging and Neuromodulation, Nathan S. Kline Institute for Psychiatric Research, Orangeburg, NY; Department of Physiology and Pharmacology, SUNY Downstate Health Sciences University, Brooklyn, NY; Department of Neuroscience, University of Wisconsin-Madison, Madison, WI, USA; Department of Neurology, Kings County Hospital, Brooklyn, NY; Department of Pharmacology, University of Virginia School of Medicine, Charlottesville, VA; Department of Psychiatry, Geisel School of Medicine at Dartmouth, Hanover, NH; Department of Biomedical Engineering, SUNY Downstate School of Graduate Studies, Brooklyn, NY; Department of Psychiatry, New Hampshire Hospital, Concord, NH; Department of Psychiatry, New York University School of Medicine, New York, NY

**Author notes:** Contributed equally (Erica Y. Griffith, Mohamed ElSayed, J. Julius Zhu).

## Abstract

Depolarizing current injections produced a rhythmic bursting of action potentials – a bursting oscillation – in a set of local interneurons in the lateral geniculate nucleus (LGN) of rats. The current dynamics underlying this firing pattern have not been determined, though this cell type constitutes an important cellular component of thalamocortical circuitry, and contributes to both pathologic and non-pathologic brain states. We thus investigated the source of the bursting oscillation using pharmacological manipulations in LGN slices *in vitro* and *in silico*. **1**. Selective blockade of calcium channel subtypes revealed that high-threshold calcium currents I_*L*_ and I_*P*_ contributed strongly to the oscillation. **2**. Increased extracellular K^+^ concentration (decreased K^+^ currents) eliminated the oscillation. **3**. Selective blockade of K^+^ channel subtypes demonstrated that the calcium-sensitive potassium current (I_*AHP*_ ) was of primary importance. A morphologically simplified, multicompartment model of the thalamic interneuron characterized the oscillation as follows: **1**. The low-threshold calcium current (I_*T*_ ) provided the strong initial burst characteristic of the oscillation. **2**. Alternating fluxes through high-threshold calcium channels and *I*_AHP_ then provided the continuing oscillation’s burst and interburst periods respectively. This interplay between I_*L*_ and I_*AHP*_ contrasts with the current dynamics underlying oscillations in thalamocortical and reticularis neurons, which primarily involve I_*T*_ and I_*H*_, or I_*T*_ and I_*AHP*_ respectively. These findings thus point to a novel electrophysiological mechanism for generating intrinsic oscillations in a major thalamic cell type. Because local interneurons can sculpt the behavior of thalamocortical circuits, these results suggest new targets for the manipulation of ascending thalamocortical network activity.

## 1 Introduction

The thalamus is important for multiple brain functions, including sensory information processing (Lörincz et al., 2008; Foxe and Snyder, 2011; Vijayan and Kopell, 2012; Lakatos et al., 2020), sleep (Bonjean et al., 2011; Coulon et al., 2012; David et al., 2013; Chen et al., 2015; Gent et al., 2018), and memory retrieval (Ketz et al., 2015). Thalamocortical oscillations are central to these operations, and aberrations in oscillatory activity have been observed in several pathological states (Zaman et al., 2011; Sitnikova et al., 2012; Avoli, 2012; Porcaro et al., 2017; Hodkinson et al., 2016).

Thalamic circuit-scale oscillations depend not only on synaptic interactions throughout the network, but also on the intrinsic electrophysiology of the involved cell types. For example, both the GABAergic cells of the thalamic reticular nucleus (TRN) and the glutamatergic thalamocortical relay (TC) cells possess intrinsic pacemaking properties that give rise to spindle and delta oscillations (Destexhe and Sejnowski, 2003; Fogerson and Huguenard, 2016). Both TRN and TC intrinsic oscillations involve low-threshold T-type calcium channels, something which has been explored extensively through both experiment and simulation (Destexhe and Sejnowski, 2003; Robinson et al., 2004; Grimbert and Faugeras, 2006; Crunelli et al., 2006; Bhattacharya et al., 2011; Wang et al., 2014; Fogerson and Huguenard, 2016).

In this study, we investigate the basis of intrinsic oscillatory firing in another major thalamic cell constituent: the local thalamic interneuron (TI). This cell type comprises up to 25%-30% of the cells in the thalamic sensory nuclei of primates, and can demonstrate tonic firing (Pape and McCormick, 1995; Halnes et al., 2011), burst firing (Zhu et al., 1999b; Halnes et al., 2011), and intrinsic oscillations (Zhu et al., 1999a; Halnes et al., 2011).

The mechanisms underlying the periodic bursting (or intrinsic oscillation) in this cell type have not yet been fully characterized. However, the behavior of these local interneurons can play an important role in sculpting thalamocortical activity (Dubin and Cleland, 1977; Wang et al., 2007; Babadi et al., 2010; Saalmann and Kastner, 2011; Wang et al., 2011b,0; Pressler and Regehr, 2013; Bastos et al., 2014; Hirsch et al., 2015; Cox and Beatty, 2017). Phasic inhibition from LGN interneurons onto thalamocortical relay cells helps sculpt the alpha rhythm observed in visual cortex (Lorincz et al., 2009). These interneurons also provide feedforward inhibition to thalamocortical relay cells in the visual thalamus (Cox and Beatty, 2017; Paz and Huguenard, 2015; Kerschensteiner and Guido, 2017; Guido, 2018). This connectivity between local thalamic interneurons and excitatory relay cells helps to sharpen feature representation in visual thalamus (Paz and Huguenard, 2015; Guido, 2018) and improve the efficiency of downstream signaling (Wang et al., 2011a; Hirsch et al., 2015). By deinactivating certain voltage-gated calcium channels, this feedforward inhibition also allows thalamocortical relay cells to switch from tonic to burst firing modes during naturalistic stimulus viewing (Wang et al., 2007). Due to the impact of local thalamic interneurons on the behavior of thalamocortical circuits, understanding the cellular mechanisms underlying this cell type’s periodic bursting behavior could prove useful for characterizing and manipulating non-pathologic and pathological states in these brain regions.

Here we have identified 39 thalamic interneurons in rat lateral geniculate nucleus (LGN) which produced sustained oscillatory bursting in response to depolarizing current injection, unchanged by synaptic blockade. We used a dual *in vitro* and *in silico* approach to delineate the contributions of specific channel conductances to the oscillation. We find that these oscillations were not due to low-threshold T-type calcium channels, as in TC and TRN cells, but instead to an interplay between high-threshold calcium conductances (I_*L*_) and a slow, calcium-activated potassium conductance (I_*AHP*_ ). These findings thus characterize a unique, newly-identified mechanism for generating intrinsic oscillations in a thalamic cell type.

## 2 Methods

Pharmacological investigation of 39 thalamic interneurons was conducted in LGN slice. Recordings were done under current clamp. Ion substitutions were performed to determine which species of ion conductances (e.g. calcium, chloride) were most critical to the intrinsic oscillation. Specific channel blockers were then used to elucidate the importance of particular channel types to the oscillation.

The data from these experiments were obtained in the 1990s and have remained unpublished. Thus, certain pharmacological blockers that might be used at present were not available at the time of these recordings – *e*.*g*., flufenamic acid to block I_CAN_ (Mrejeru et al., 2011).

Physiological experiments were complemented by computer simulation of a model interneuron. A cell model with simplified morphology was used, with soma and dendrite geometry set to roughly match the dimensions observed in these interneurons. Channel types and parameters were informed by the literature (Augustinaite et al., 2011; Acuna-Goycolea et al., 2008; Allken et al., 2014; Zhu and Lo, 1999; Crunelli et al., 1988; Munsch et al., 1997; Pape and McCormick, 1995; Pape et al., 1994) and by previously obtained voltage clamp results (Zhu et al., 1999b,9). Additional ion channel species were inserted into the model to incorporate the *in vitro* electrophysiology data from the ion substitution and pharmacological blockade experiments. The model was then used to further examine the channel dynamics underlying the intrinsic oscillation. Code and data regarding both the model and the physiological experiments have been deposited on Mendeley at https://data.mendeley.com/datasets/y7czc2hckf/draft?a=5deb096e-6621-48b7-831e-cc029cc1a741.

### 2.1 Electrophysiology

Slice preparation was described previously (Zhu and Uhlrich, 1997; Zhu et al., 1999a) and approved by the Institutional Animal Care and Use Committee of the University of Wisconsin School of Medicine and Public Health. Sprague-Dawley rats (100-300g) were deeply anesthetized by halothane. The brain was quickly removed after decapitation and placed into cold (6 – 8°C) physiological solution containing NaCl (126 mM), KCl (2.5 mM), NaH_2_PO_4_ (1.25 mM), NaHCO_3_ (26 mM), MgSO_4_ (1 mM), dextrose (20 mM), CaCl_2_ (2 mM), at pH 7.35. The solution was continuously bubbled with 95% O_2_-5% CO_2_. Slices containing LGN, each 500mm thick, were cut from the tissue blocks with a microslicer. Slices were kept in oxygenated physiological solution for a minimum of 2 hours before recording. Methods for tight-seal patch recordings also match those described in previous papers (Zhu et al., 1999c,9,9). Patch electrodes of 7-9 MΩ resistance were used in whole-cell recording configuration, with a standard intracellular solution consisting of: C_6_H_11_O_7_K (120 mM), N-2-hydroxyethylpiperazin-N’-2-ethanesulphonic acid (HEPES) (10 mM), ethyleneglycolbis(amino-ethylether)tetra-acetate (EGTA) (5 mM), MgCl_2_ (2 mM), adenosine triphosphate (4 mM), guanosine triphosphate (0.1 mM), CaCl_2_ (0.5 mM), KCl (10 mM), and biocytin 0.25%, at pH 7.25.

Recordings were done with an Axoclamp-2A amplifier (Axon Instruments). The chamber was perfused with oxygenated physiological solution comprised of: NaCl (126 mM), KCl (2.5 mM), NaH_2_PO_4_ (1.25 mM), NaHCO_3_ (26 mM), MgSO_4_ (1 mM), dextrose (20 mM), CaCl_2_ (2 mM), at pH 7.35. Half time for the bath solution exchange was approximately 7 seconds. The bath temperature was kept at 34.0 *±* 0.5^*?*^C. Blockers were applied with the bath solution. The specific pharmacological agents used are detailed in the text. All chemicals were purchased from Sigma. Recordings routinely lasted 1-6 hours and were generally stable, showing little change in input resistance and resting membrane potential (-67.7*±* 5.1 mV) throughout the recording period.

Interneurons were identified physiologically using established criteria (Munsch et al., 1997; Pape and McCormick, 1995; Zhu and Uhlrich, 1997). Interneurons had a higher input resistance (*>* 350 MΩ) compared to thalamocortical cells (*<* 200 MΩ) (Guido, 2018; Zhu et al., 1999b), as well as a longer membrane time constant (*>* 50 ms) compared to thalamocortical cells (*<* 40 ms) (Zhu et al., 1999a). These two properties were sufficient to distinguish local thalamic interneurons from other thalamic cell types (Zhu et al., 1999b). The identity of all interneurons was also confirmed morphologically with histological visualization, using the avidin-biotin-peroxidase method (Horikawa and Armstrong, 1988).

### 2.2 Simulation

Cell models were instantiated using the NEURON simulator (Hines, 1993) and adapted for use in NetPyNE (Dura-Bernal et al., 2019). Model interneurons consisted of 15 cylindrical compartments: a soma, 2 equivalent proximal dendrites, and 12 equivalent distal dendrites. Passive parameters (e.g. leak channel density) were tuned to replicate the charging curve (Zhu et al., 1999b,9). Basic spiking behavior was produced using Hodgkin-Huxley type sodium and potassium channels. A hyperpolarization-activated cation current (I_H_) (Zhu et al., 1999c), a calcium-sensitive inward current (I_CAN_) (Zhu et al., 1999b), and a low-threshold calcium current (I_T_) (Zhu et al., 1999b) were also included based on previously modeled voltage clamp results (Zhu et al., 1999c,9). A high-threshold calcium current (I_L_) and a slow, calcium-dependent potassium current (I_AHP_) were added to the model when pharmacological blockade results suggested their involvement in the intrinsic oscillation. Channel mechanisms for I_L_ and I_AHP_ currents were modified from existing parameterizations which utilized data from different animal models (e.g. mouse) or different thalamic cell types (e.g. thalamic reticular cells) (Destexhe et al., 1994a; Halnes et al., 2011; McCormick and Huguenard, 1992). These channels were added in a bootstrap manner (Marder and Abbott, 1995), in tandem with pharmacological investigation.

Diameters (D) and lengths (L) of the cylindrical compartments were estimated from stained geniculate interneurons and adjusted to fit the membrane time constant obtained by small current injections. Passive parameters and geometry are listed in Table 1. Note that small variations in the maximum conductance or reversal potential of the leak channels can be used to capture the natural variation observed *in vitro*, in properties such as the resting membrane potential.

**Table 1:** Geometry and passive parameters for the 15-compartment cell model. D is the diameter and L is the length of each compartment. C_m_ is the capacitance, R_a_ the axial resistivity, 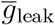 the maximum conductance of the leak channel, and E_leak_ is the reversal potential of the leak channel.

| Section | D ($\mu\text{m}$) | L ($\mu\text{m}$) | $C_m$ ($\frac{\mu\text{F}}{\text{cm}^2}$) | $R_a$ ($\Omega\cdot\text{cm}$) | $\bar{g}_{\text{leak}}$ ($\frac{\text{mS}}{\text{cm}^2}$) | $E_{\text{leak}}$ (mV) |
| --- | --- | --- | --- | --- | --- | --- |
| Soma | 10 | 16 | 1 | 100 | 0.025 | -72 |
| Proximal Dendrite | 3.25 | 240 |  |  |  |  |
| Distal Dendrite | 1.75 | 180 |  |  |  |  |

The Hodgkin-Huxley parallel conductance model was used for active channels. For the results shown here, active channels were inserted exclusively into the soma compartment of the model cell. We used the Borg-Graham modification (Borg-Graham, 1991) of the Hodgkin-Huxley parameterizations for the I_Naf_ (fast sodium) and I_Kdr_ (delayed rectifier) currents (Lytton and Sejnowski, 1991; Traub et al., 1991). Parameterizations for the I_H_, I_T_, I_L_, I_CAN_, and I_AHP_ channel mechanisms are described below. For these channel mechanisms, the change in a given state variable (e.g. m, h) with respect to time can be described by:

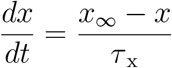

Parameters for I_H_ (a hyperpolarization-activated cation current) were obtained from previously gathered voltage-clamp data (Zhu et al., 1999c). The reversal potential for this channel mechanism was E_H_ = -44 mV. The current can be described by following parameterization:

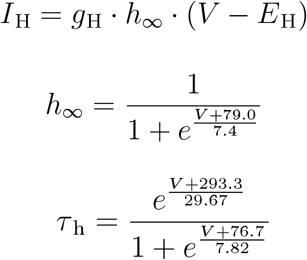

Values for I_T_ (a low-threshold calcium channel) were also obtained from previous voltage-clamp data (Zhu et al., 1999b). Kinetics were corrected for temperature using m_Q10_=3 and h_Q10_=1.5. This channel mechanism employed the Goldman-Hodgkin-Katz formalism:

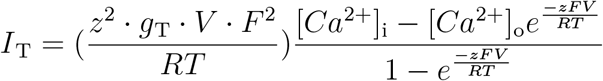

where z=2 (for Ca^2+^), F represented the Faraday constant, T represented the temperature in Kelvin, R represented the Boltzmann gas constant, and internal calcium concentration [Ca^2+^]_i_ = 50 nM and external calcium concentration [Ca^2+^]_o_ = 2 nM. Conductance g_T_ was defined by:

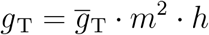

with 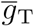 representing the maximum possible conductance value for this channel, and with state variables m and h (and their corresponding time constants) described by the following equations:

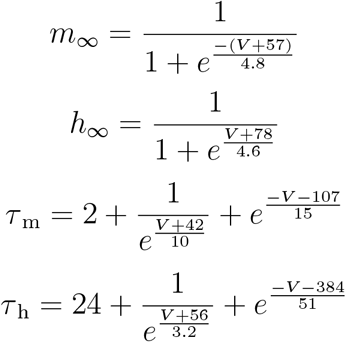

I_L_ (a high-threshold calcium current) was adapted from a previous study (Kay and Wong, 1987) with kinetics corrected for temperature using Q_10_ = 3. This channel mechanism also used the Goldman-Hodgkin-Katz formalism, as described above for I_T_. Conductance for this channel was defined by:

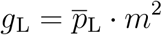

with 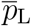 representing the maximum permeability of the channel. The activation variable m and corresponding time constant *τ* _m_ were described by:

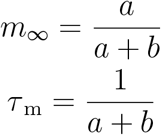

where

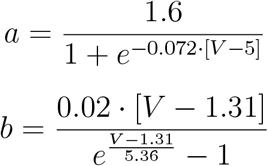

Two separate calcium pools were simulated, with one corresponding to influx from the low-threshold calcium conductance (I_T_) and the other corresponding to influx from the high-threshold calcium conductance (I_L_). This was done due to prior results (Zhu et al., 1999b) suggesting that the I_CAN_ and I_AHP_ currents may have different sensitivities to calcium ions from high-vs. low-threshold calcium channels. The two calcium pumps used for these pools were identical, with separate time constants for high and low Ca^2+^ concentrations:

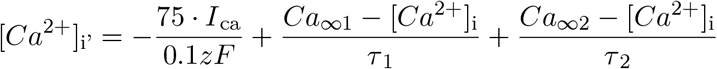

where the low steady-state calcium concentration Ca_*∞*1_ = 0.01 nM, and the high steady-state calcium concentration Ca_*∞*2_ = 52 nM, and time constants *τ* _1_=150 ms and *τ* _2_ = 80 ms.

I_CAN_ (a calcium-activated cation current) was adapted from a previous study (Destexhe et al., 1994b) and modified to fi t ou r voltage-clamp da ta (Z hu et al., 1999b), with reversal potential E_CAN_ = 10 mV and the following parameterization describing the current:

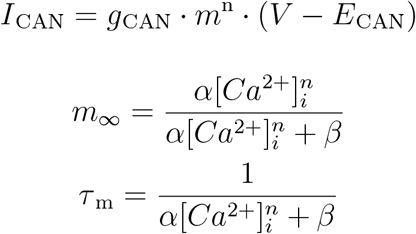

where *β*=0.003 ms^-1^, *α*=1.4e5 ms^-1^*µ*M^-8^, and n=8.

I_AHP_ (a calcium-activated potassium current) used the same parameterization as I_CAN_ (above), but with E_AHP_ = -95 mV, *β*=0.02 ms^-1^, *α*=0.031 ms^-1^*µ*M^-2^, and n=2.

## 3 Results

### 3.1 Calcium Conductances

Local thalamic interneurons possess both high-(Acuna-Goycolea et al., 2008; Budde et al., 1998) and low-threshold (Parajuli et al., 2010; Allken et al., 2014) calcium conductances. Reducing the calcium driving force also prevented burst firing in LGN interneurons in a prior study (Zhu et al., 1999b). Calcium conductances underlie bursting and intrinsic oscillations in both thalamocortical relay cells (Zhang et al., 2002; Destexhe and Sejnowski, 2003; Crunelli et al., 2006; Dilger et al., 2011; Kim et al., 2015; Fogerson and Huguenard, 2016) and thalamic reticular cells (Destexhe and Sejnowski, 2003; Chausson et al., 2013; Coulon et al., 2009; Crandall et al., 2010; Fogerson and Huguenard, 2016). We thus used selective calcium channel blockades to determine which calcium currents contributed to the bursting oscillation in this cell type.

#### 3.1.1 High-threshold Calcium Conductances: *in vitro*

Generalized high-threshold calcium channel blockade can be achieved with Cd^2+^ or Co^2+^ (Swandulla and Armstrong, 1989; Lansman et al., 1986; Chow, 1991; Ryu and Randic, 1990). Bath application of Cd^2+^ (200 *µ*m; n=4) (Swandulla and Armstrong, 1989; Lansman et al., 1986; Chow, 1991; Shen et al., 2000) reversibly blocked the oscillation (Fig. 1B). The oscillation recovered with removal of Cd^2+^ (Fig. 1C). Application of Co^2+^ (1 mM; n=2) (Ryu and Randic, 1990) achieved similar results (Fig. 1D). Recovery was not achieved in these cells before recordings were lost.

**Figure 1:**
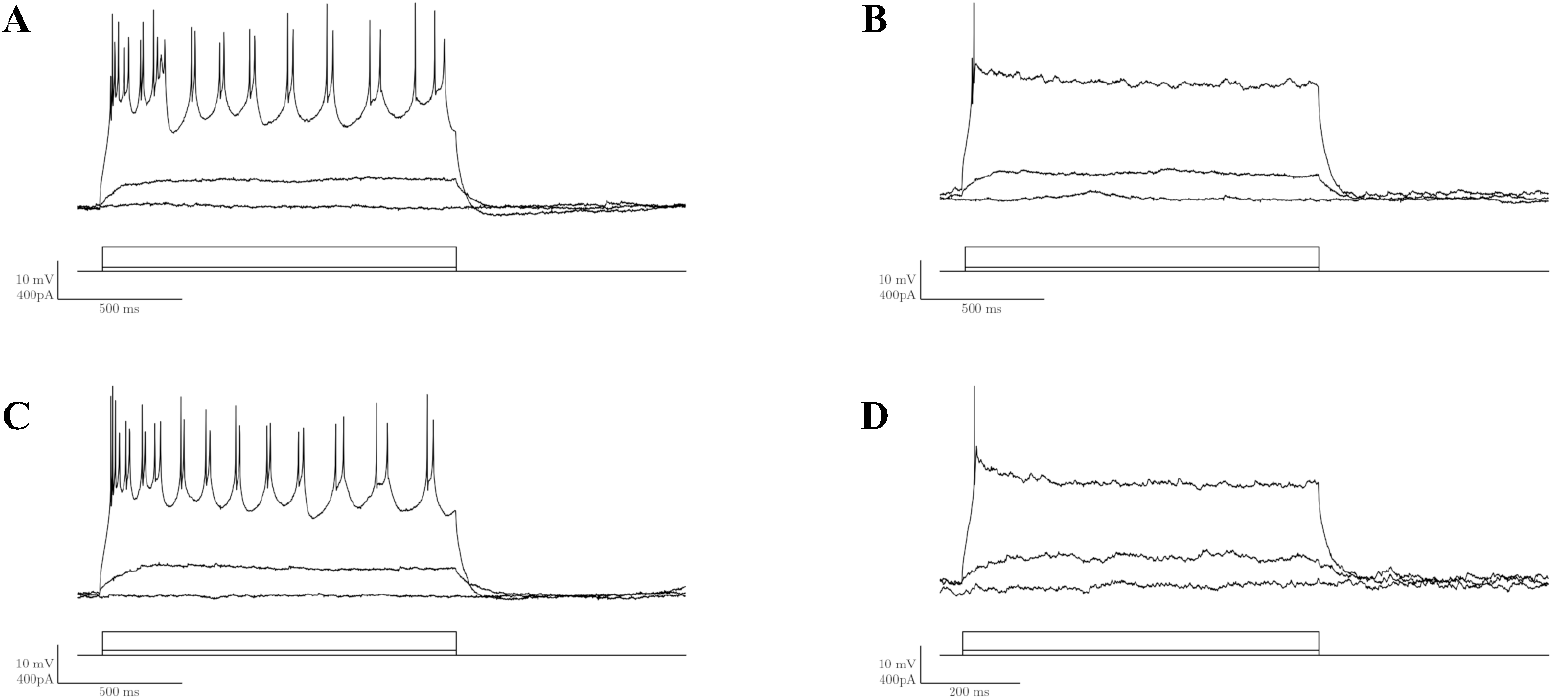
Generalized blockade of high-threshold calcium conductances abolishes bursting oscillations *in vitro*. A: Depolarizing current injection induced bursting oscillation. B: Bath application of a high-threshold calcium conductance blocker, Cd^2+^ (200*µ*M), blocked the oscillation. C: After washout of Cd^2+^, the oscillation was recovered. D: Subsequent to C, addition of Co^2+^ (1 mM), another high-threshold calcium conductance blocker, also blocked the oscillation.

Selective blockades of the N-, Q-, L-, and P-type calcium channels were performed next. Blockades of N- and Q-type calcium channels showed little to no effect on the oscillation. Neither *ω*-conotoxin GVIA (5 *µ*M, n=3), a highly-specific N-type calcium channel blocker (Adams and Berecki, 2013; Nielsen et al., 2000; McDonough et al., 2002, 1996; Randall and Tsien, 1995; Wheeler et al., 1994), nor *ω*-conotoxin MVIIC (2 *µ*M, n=2), an N- and Q-type channel blocker (Christina I. Schroeder, 2006; McDonough et al., 1996; Wheeler et al., 1994), impaired the oscillation.

In contrast, L-type channel blockade, achieved with bath application of 20 *µ*M nifedipine (Shen et al., 2000; Lambert and Wilson, 1996; Wisgirda and Dryer, 1994), enhanced the initial burst and increased the oscillation frequency (n=4; Fig. 2B). Similar results were obtained with nimodipine, another L-type calcium channel blocker (Randall and Tsien, 1995; Wheeler et al., 1994). P-type calcium channel blockade also resulted in an enhanced initial burst followed by increased oscillation frequency (n=3; Fig. 2E). P-type blockade was achieved using 50 *µ*M *ω*-conotoxin MVIIC (McDonough et al., 2002, 1996; Nimmrich and Gross, 2012; Wheeler et al., 1994); the effect was not reversed after washing with normal bath solution for up to 35 minutes.

**Figure 2:**
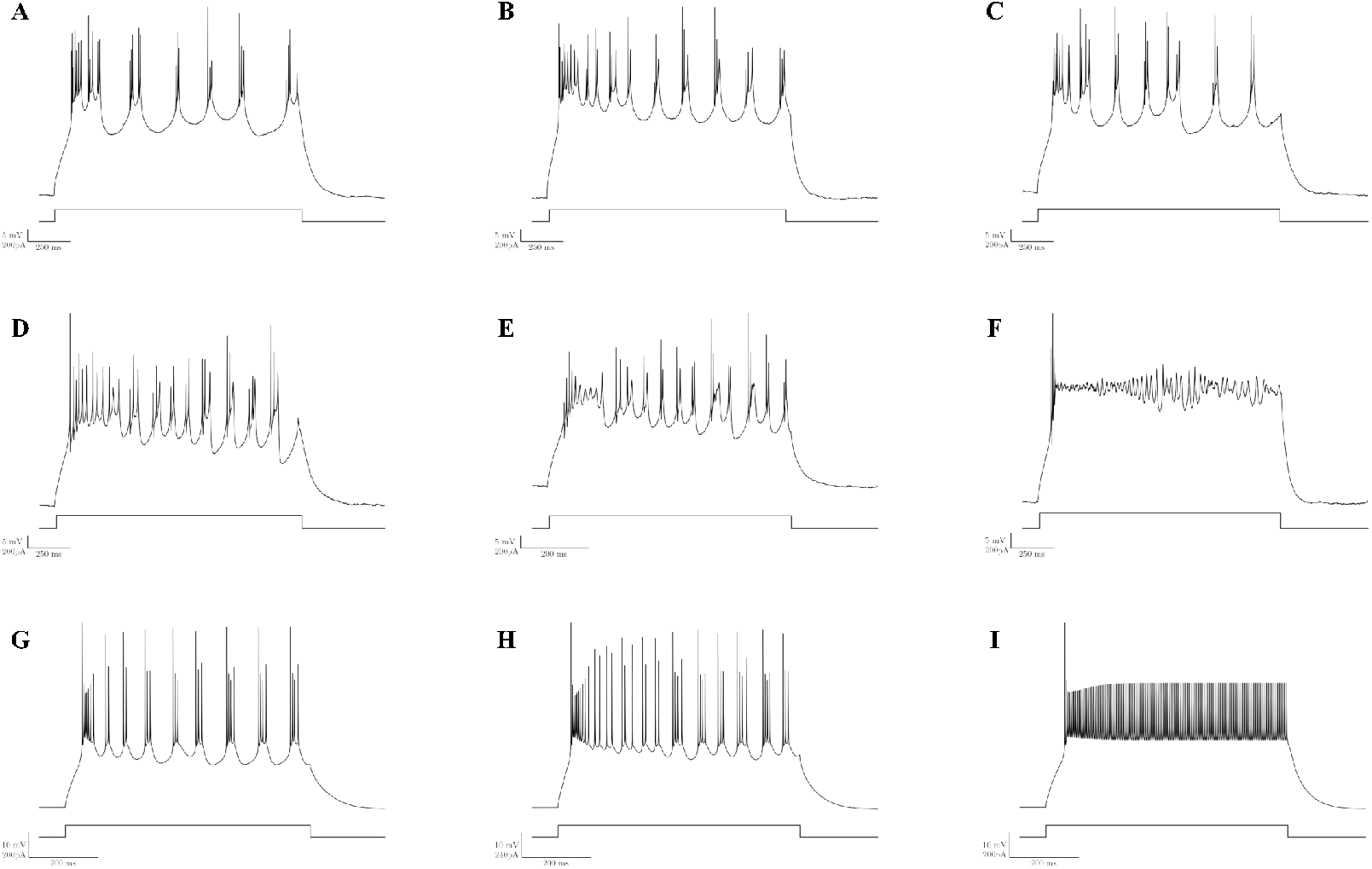
Combined blockade of specific high-threshold calcium channels blocks the oscillation. A-F: High-threshold calcium-channel manipulations *in vitro*. G-I (bottom row): *in silico* calcium channel manipulations. A: Control, *in vitro*. B: Selective L-channel blockade (20 *µ*M nifedipine) enhances the initial burst and subsequent oscillation rate *in vitro*. C: Nifedipine washout. D: Control, *in vitro*. E: Selective P-channel blockade (50 *µ*M MVIIC) alters the initial burst and increases oscillation rate *in vitro*. F: Combined L- and P-type calcium channel blockade (20 *µ*M nifedipine + 50 *µ*M MVIIC) abolishes the oscillation. G: Control, *in silico*. H: Reduction of high-threshold calcium channel conductance to approximately ⅔of the control value enhances the initial burst and increases the oscillation rate *in silico*. I: Reduction of high-threshold calcium channel conductance to approximately ⅓of the control value abolishes the *in silico* oscillation.

Generalized and selective calcium channel blockades thus produced different effects. However, simultaneous blockade of L- and P-type channels using nifedipine (20 *µ*M) and *ω*-conotoxin MVIIC (50 *µ*M) resulted in an initial spike followed by a persistently depolarized state (Fig. 2F). Qualitatively similar traces resulted from the generalized high-threshold calcium blockades (Fig. 1B, D). From this, we concluded that L- and P-type calcium channels together may provide the bulk of the high-threshold calcium current responsible for the oscillation dynamics.

#### 3.1.2 High-threshold calcium conductances: *in silico*

We modeled the high-threshold calcium currents as a single conductance (I_L_) *in silico*. Decreasing the density of the I_L_ channels to approximately 2*/*3 of the control value resulted in an enhanced initial burst and an increased oscillation frequency (Fig. 2H). A more drastic decrease in I_L_ channel density, to approximately 1*/*3 of the control value, resulted in an initial spike followed by a persistently depolarized state (Fig. 2I). These changes aligned qualitatively with our *in vitro* results (Fig. 2F), lending credence to the validity of the *in silico* results.

#### 3.1.3 Low-threshold calcium conductances

Next we assessed the role of low-threshold calcium conductances in the generation and maintenance of the oscillation. At concentrations of 50 *µ*M, Ni^+^ has been shown to block low-threshold T-type calcium channels (Kang et al., 2006; Lee et al., 1999; Nikonenko et al., 2005). However, at higher concentrations (e.g. 100 *µ*M), Ni^+^ has been found to alter current flow through high-threshold calcium channels as well, by both blocking current flow and shifting the voltage required for activation (Zamponi et al., 1996; Lee et al., 1999; McFarlane and Gilly, 1998). In general, concentrations of 50 *µ*M Ni^+^ did not substantially affect the oscillation, instead variably diminishing or delaying the initial burst of the oscillation. This effect was mimicked by our *in silico* observations (Fig. 3), where even a 40% reduction in I_T_ channel conductance resulted in a delayed initial spike.

**Figure 3:**
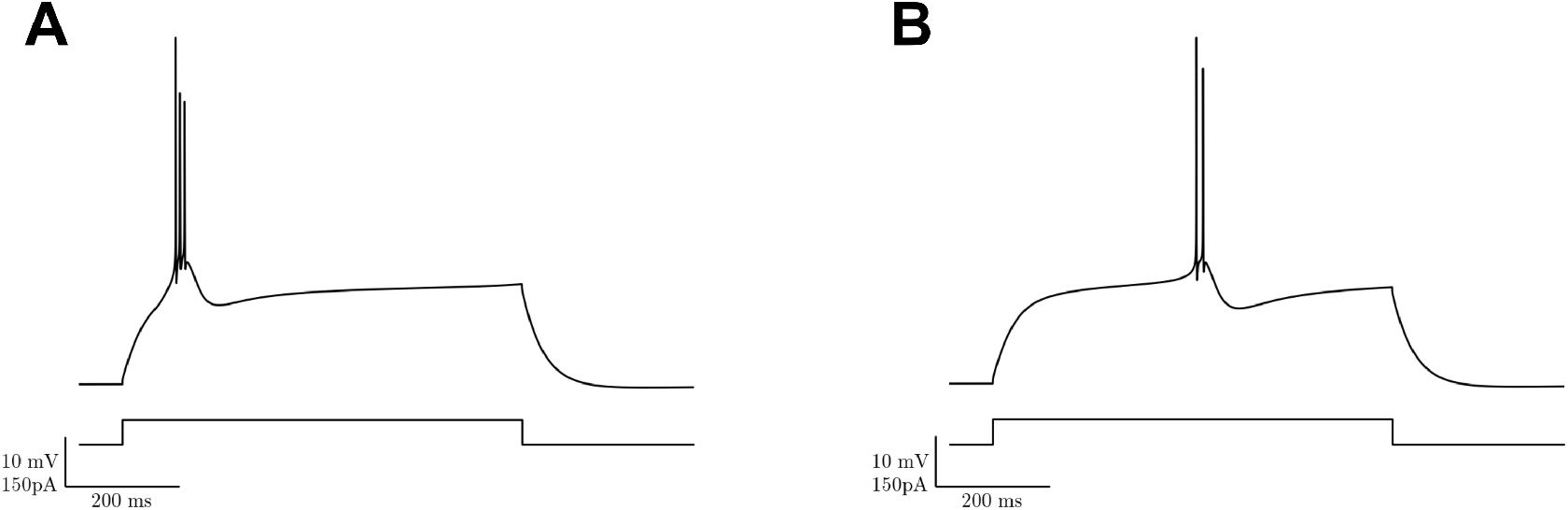
Low-threshold calcium-channel manipulation, *in silico*. A: Control. B: Voltage trace after 40% reduction in low-threshold calcium channel conductance.

### 3.2 Depolarizing Cation Conductances

#### 3.2.1 I_H_ Conductance

We have previously demonstrated that local interneurons in LGN have a prominent I_H_ current (Zhu et al., 1999c). This current is described as an anomalously-rectifying inward current with a mixed sodium and potassium conductance. We suspected that the oscillation would not involve I_H_, as the relevant channels would be largely inactivated at the level of depolarization seen during the oscillations (Zhu et al., 1999c). Bath application of Cs^2+^, an I_H_ blocker (Thoby-Brisson et al., 2000; Bonin et al., 2013; Kitayama et al., 2003; Svoboda and Lupica, 1998; Banks et al., 1993; Galligan et al., 1990), showed little effect on the oscillation. Simulated blockade of I_H_ conductance *in silico* also demonstrated little effect.

#### 3.2.2 I_CAN_ Conductance

We previously identified a plateau potential in local geniculate interneurons that was most likely mediated by I_CAN_ (Zhu et al., 1999b), a calcium-activated nonselective cation current. We used the model to investigate the activity of each conductance throughout the oscillation, and observed that I_CAN_ contributes depolarizing current in the period of time prior to each burst (Fig. 11). We then decreased I_CAN_ conductance *in silico*, and found that elimination of this conductance did not abolish the oscillation. However, consistent with our first observation (namely that I_CAN_ helps to depolarize the cell prior to bursting), elimination of I_CAN_ both reduced the oscillation frequency and increased the time to first burst (Fig. 4). Thus, although I_CAN_ appears to facilitate and support the oscillation, we predict that this conductance is not crucial to its formation.

**Figure 4:**
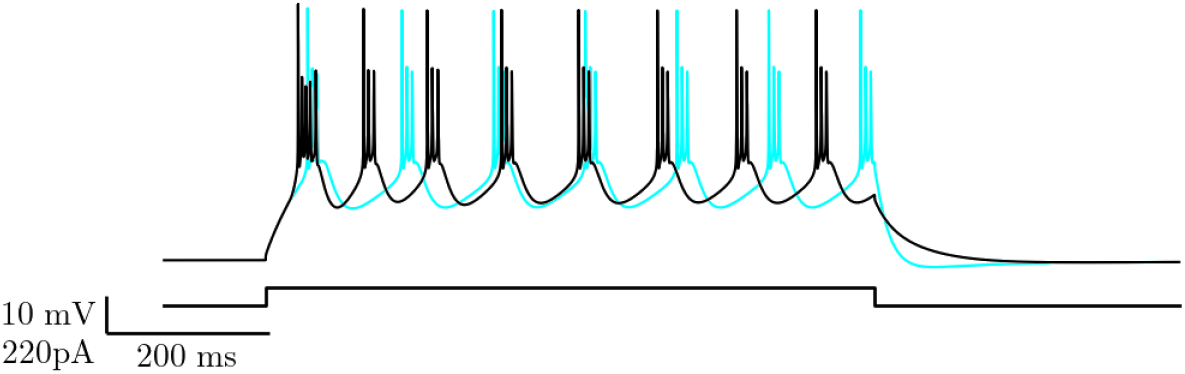
Elimination of the I_CAN_ conductance *in silico* increased the time to initial burst and reduced the overall oscillation frequency. Black: Control I_CAN_ density. Cyan: I_CAN_ density set to 0.

#### 3.2.3 I_Naf_ Conductance

Tetrodotoxin (TTX), the voltage-gated sodium channel blocker, eliminated the oscillation *in vitro*, while while synaptic blockade by receptor antagonists did not (Zhu et al., 1999a). This result suggested that the oscillation arose as the result of intrinsic cell properties, and also suggested a role for fast sodium channels in the maintenance of the oscillation. We used the model to manipulate I_Naf_ current *in silico* to investigate this further. Reducing the density of the fast sodium channels by 75% resulted in lower spike amplitudes, while the oscillation itself was maintained (Fig. 5B). Full blockade of I_Naf_ eliminated both the fast spikes and the underlying slow oscillation (Fig. 5C), matching the previously found *in vitro* results.

**Figure 5:**
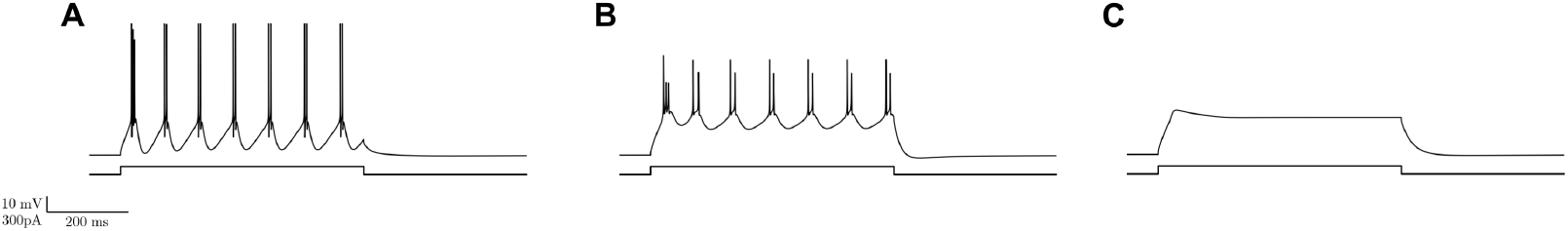
Effect of reducing I_Naf_ density *in silico*. A: Control. B: 75% reduction reduces amplitude of spikes but maintains oscillation. C: Elimination of I_Naf_ eliminates oscillation.

### 3.3 Hyperpolarizing Conductances

In order for oscillation to occur, hyperpolarizing driving forces must be present to sculpt the interburst interval. Chloride and potassium conductances were considered the two main candidates that could provide the hyperpolarization necessary to produce this part of the oscillation. Reduction of chloride driving force was achieved by replacing Cl^-^with isethionate ions, reducing extracellular chloride concentration to as low as 4 mM. This intervention had no appreciable effect on the oscillation. In contrast, increasing the extracellular K^+^ concentration to 9 mM (control value: 2.5 mM) eliminated the oscillation, with the cells instead demonstrating a depolarized plateau response (Fig. 6C). We thus focused our investigation on the role of potassium conductances.

**Figure 6:**
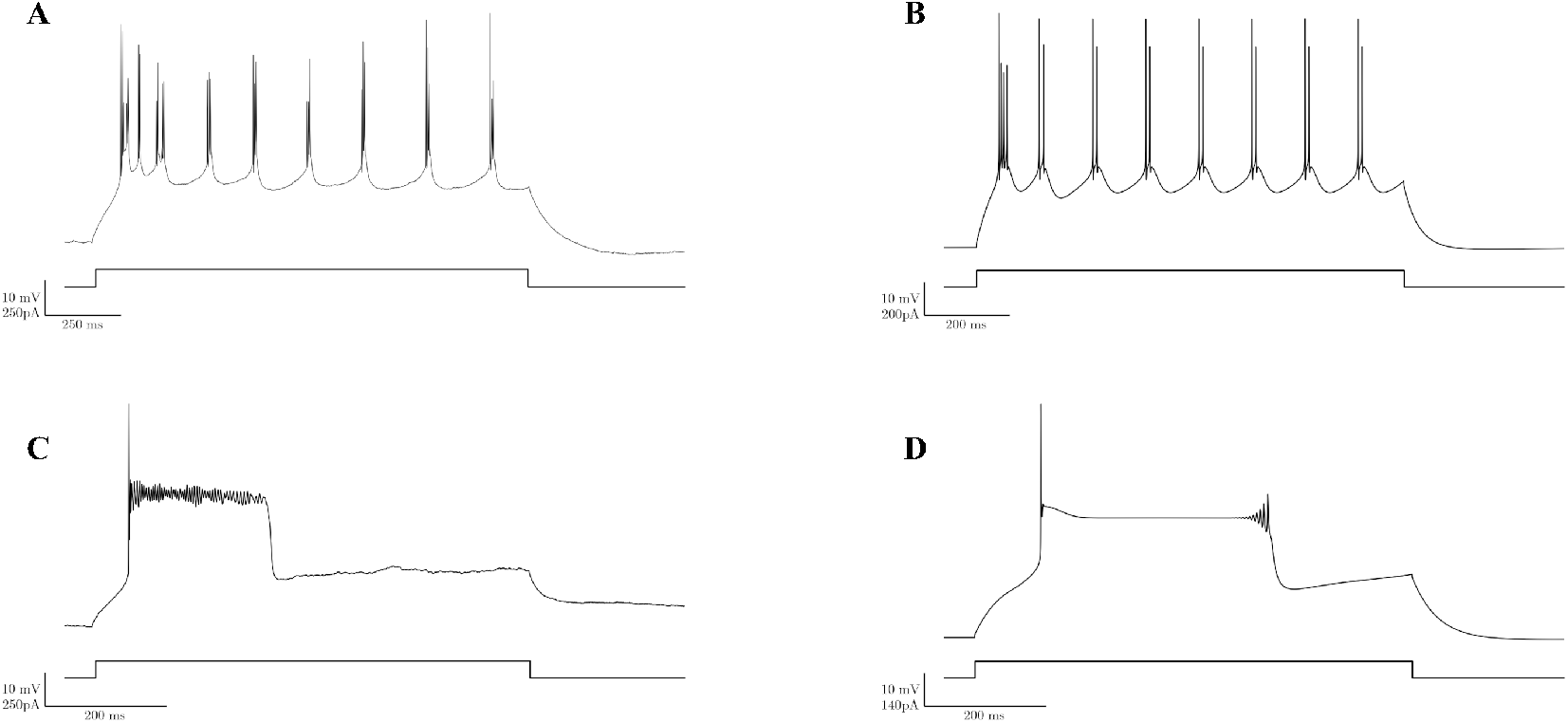
Increased extracellular potassium concentration eliminated the oscillation, resulting in a depolarized plateau potential of limited duration. Left (A, C): *In vitro* voltage traces. Right (B, D): *In silico* voltage traces. A: Control (2.5 mM K^+^), *in vitro*. B: Control, *in silico*. C: Increased extracellular [K^+^] (9 mM), *in vitro*. D: Simulated increase in extracellular [K^+^], *in silico*.

#### 3.3.1 Potassium Conductances

We examined the potential role of I_C_, a large-conductance calcium- and voltage-dependent fast potassium current. Charybdotoxin (CTX) acts as a selective blocker of the I_C_ current (Goh et al., 1992; Garcia et al., 1995). Application of 60 nM CTX did not show an appreciable effect on the oscillation (n=4; Fig. 7).

**Figure 7:**
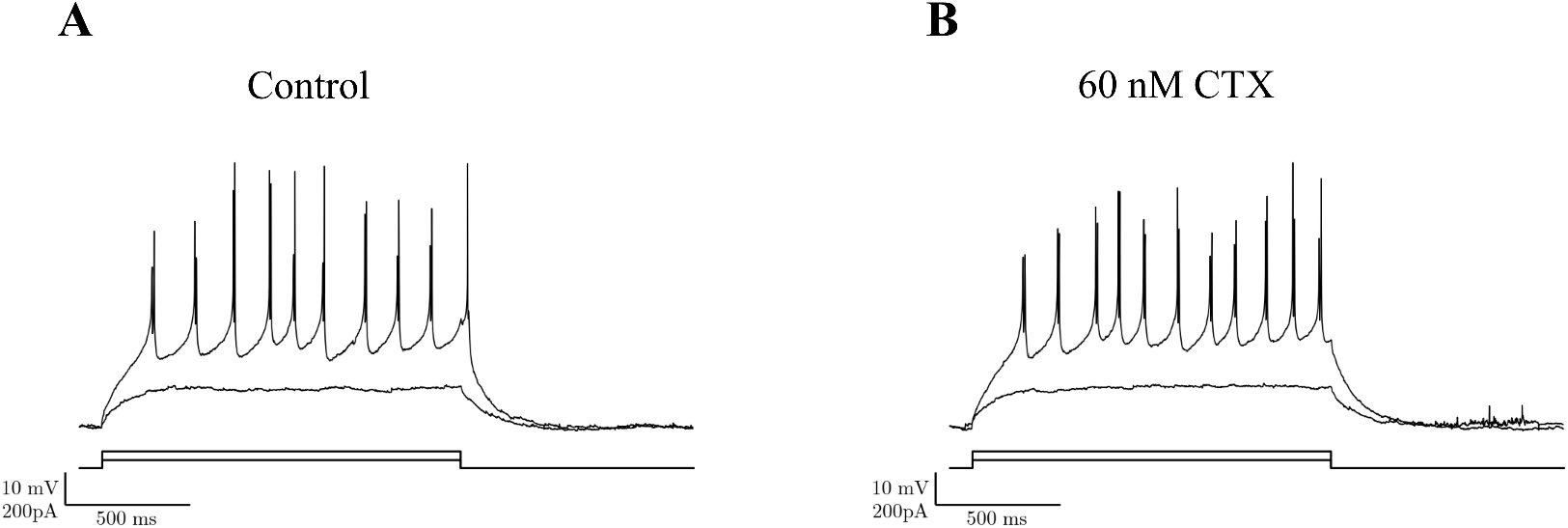
I_C_ current blockade did not impact the oscillation. A. Depolarizing current injection induced bursting oscillation. B. Application of 60 nM CTX had little effect on the oscillation (n=4).

The fast, 4-AP sensitive A-type potassium current (I_A_) has been reported to suppress the I_T_-mediated low-threshold spike in dissociated geniculate interneurons (Pape et al., 1994), making it an important target of investigation. Here, bath application of 400*µ*M 4-AP created a square-wave firing pattern (n=2; Fig. 8B) (Zhu, 1996). This effect did not reverse after a prolonged wash with normal bath solution. Although application of 4-AP prevented full interspike repolarization during the burst, it did not significantly affect the oscillatory rhythm itself. Thus, we concluded that the I_A_ current present in these interneurons activates relatively rapidly and is involved in high-frequency spikes, and not in low-frequency repetitive bursts. This conclusion aligns with the role of I_A_ in other cell types, where it has been implicated in rapid spike repolarization in pyramidal cells (Pathak et al., 2016) and hippocampal interneurons Zhang and McBain (1995). Because these two currents (I_A_ and I_C_) did not show an appreciable effect on the oscillation *in vitro*, they were not included in our simulations.

**Figure 8:**
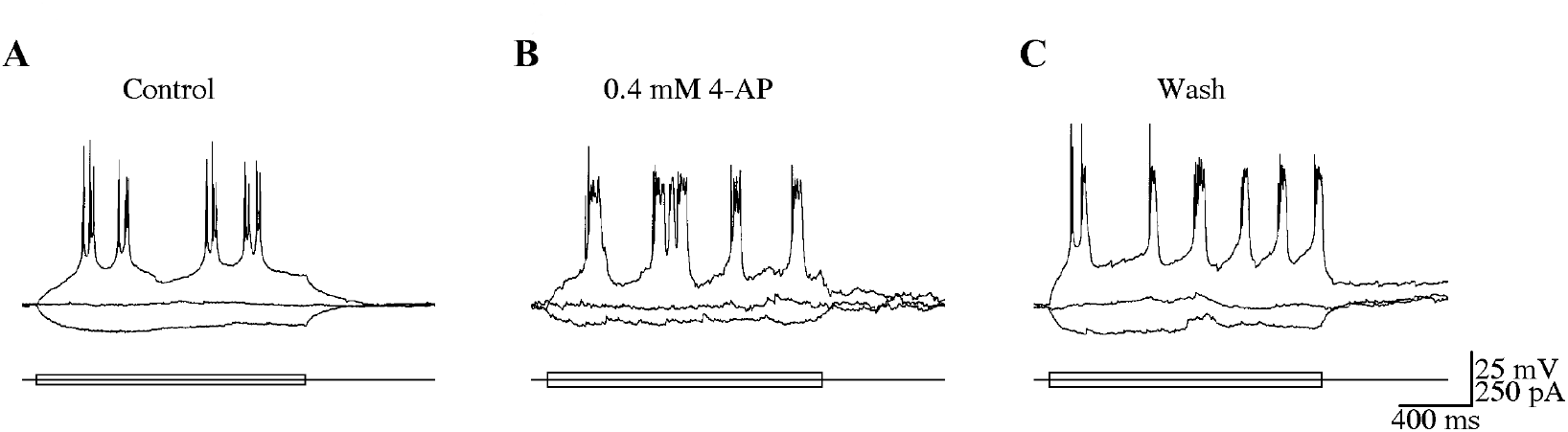
Blockade of I_A_ current prevented fast intraburst repolarization without impacting oscillation frequency. A: Depolarizing current injection induced bursting oscillation *in vitro*. B: Application of 400*µ*M 4-AP created a square-wave firing pattern, eliminating intraburst action potentials but having little impact on the oscillation frequency (n=2). C: Washout did not recover original firing pattern.

Lastly, we explored a calcium-activated potassium current, I_AHP_, that has been implicated in the oscillatory dynamics of thalamic reticular cells (Destexhe and Sejnowski, 2003; Destexhe et al., 1994a). This current also exhibits kinetics compatible with the duration of the interburst hyperpolarization (Gurney et al., 1987; Schwindt et al., 1988). To test for the presence of this current in local geniculate interneurons, we applied apamin, a specific I_AHP_ blocker (Matthews and Lee, 1991; Oh et al., 2000; Bond et al., 2004). Application of 200 nM apamin eliminated the oscillation while preserving fast spiking (n=4; Fig. 9C). The effect of I_AHP_ blockade in the model was similar, with *in silico* I_AHP_ blockade showing prolonged high-frequency spiking without oscillation (Fig. 9D). We thus concluded that the I_AHP_ current provided the interburst hyperpolarization necessary for the oscillation.

**Figure 9:**
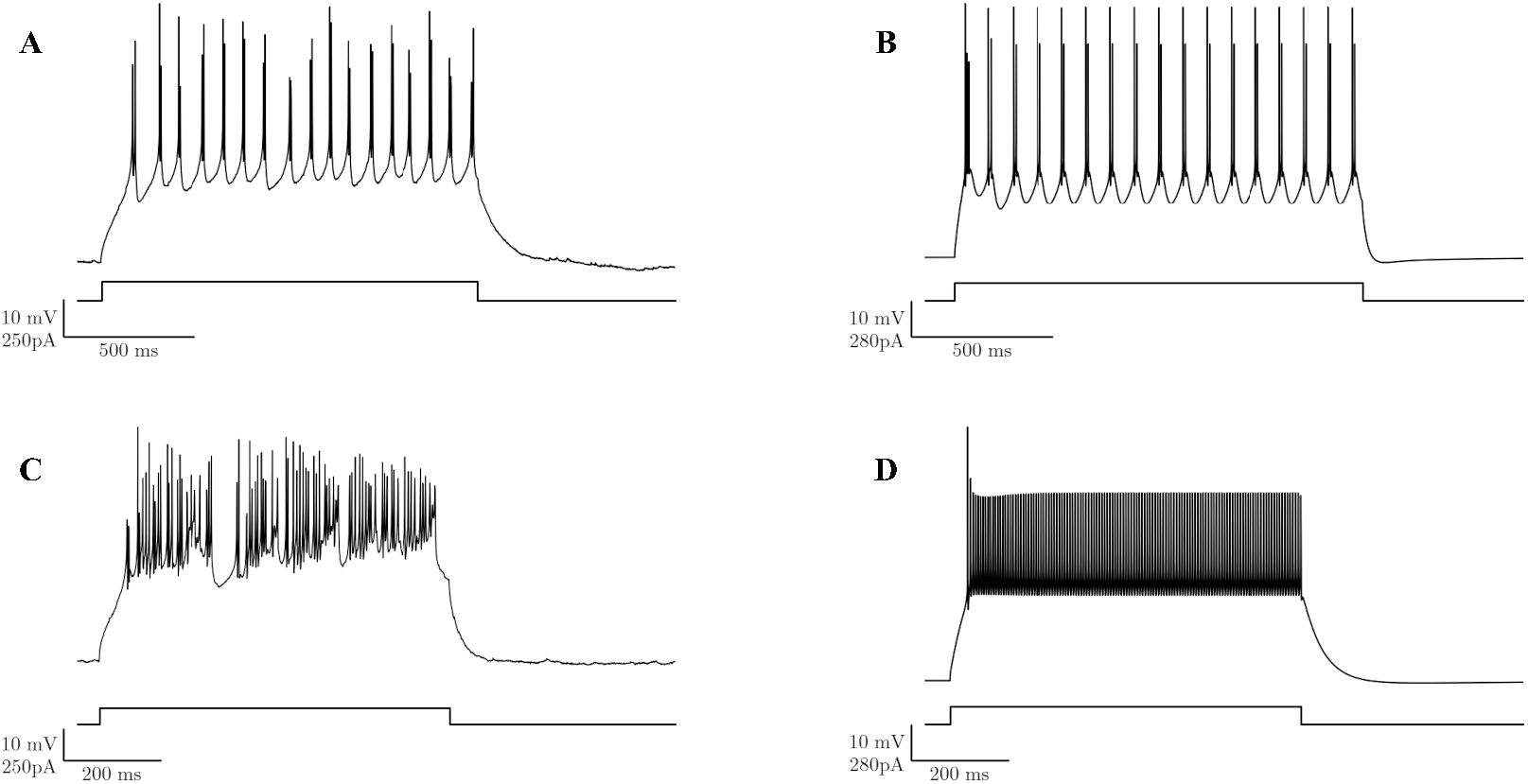
I_AHP_ contributes to the bursting oscillation. Left (A, C): Voltage traces, *in vitro*. Right (B, D): Voltage traces, *in silico*. A: Control, *in vitro*. B: Control, *in silico*. C: *In vitro* application of the I_AHP_ blocker apamin eliminated the bursting oscillation, leading to low-amplitude, high-frequency spiking. D: Elimination of the I_AHP_ conductance *in silico* similarly resulted in low-amplitude, high-frequency spiking.

### 3.4 Input Resistance

Local thalamic interneurons were consistently noted to have a higher input resistance than thalamocortical relay cells (Zhu et al., 1999a,9). Previous observations found that cells which underwent a substantial loss of input resistance (*e*.*g*., via rupture of the gigaseal during recording) were subsequently unable to sustain bursting oscillations. These cells transitioned to a single-spike tonic firing pattern in response to depolarizing current injections (Zhu et al., 1999a). We simulated this here by increasing the conductance of the I_leak_ channel. This change induced a shift to tonic firing in the cell model (Fig. 10A,B).

**Figure 10:**
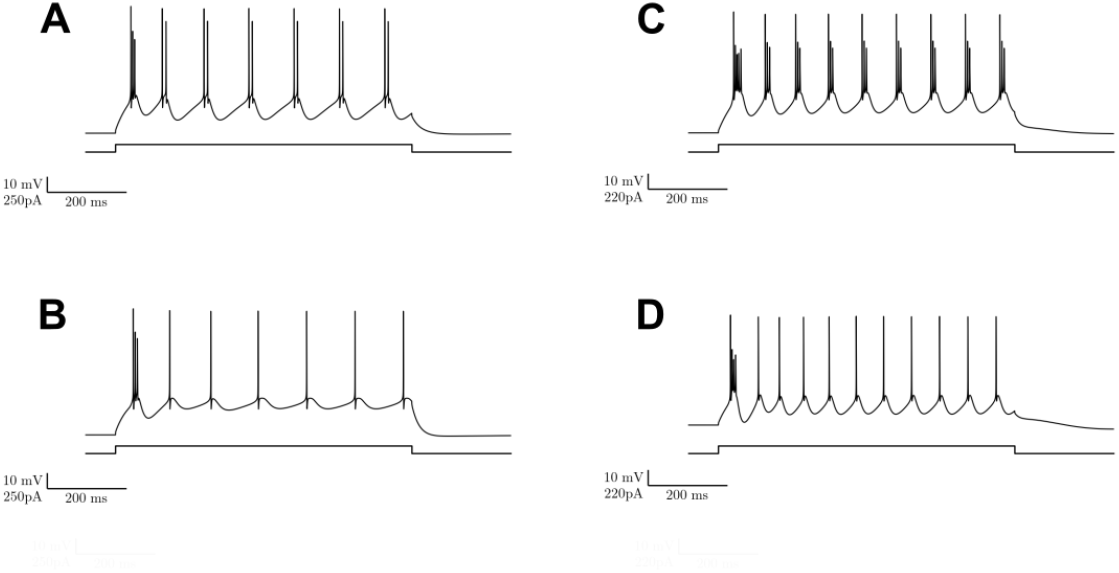
Reduced input resistance was simulated via two different *in silico* mechanisms, resulting in a shift in firing mode in both cases, from bursting oscillation to tonic firing. A: Control. B: Increased leak channel conductance. C: Control. D: Increased I_H_, I_CAN_, and I_AHP_ conductances.

It has also been noted that activation of muscarinic receptors, via application of agonists such as acetyl-*β*-methcholine, can switch the firing pattern in thalamic interneurons from bursting oscillation to tonic firing (Zhu and Heggelund, 2001). This has been attributed again to a decrease in input resistance, in this case mediated by increased conductance of I_H_ and I_CAN_ currents, along with increased conductance of a potassium current (Zhu and Heggelund, 2001; McCormick and Pape, 1988; Zhu and Uhlrich, 1998,9). We tested this *in silico*, by increasing the I_H_, I_CAN_, and I_AHP_ conductances. This also resulted in a switch from burst oscillation to tonic firing (Fig. 10C,D).

### 3.5 Interaction of Currents

Using pharmacological blockade and *in silico* manipulation, we demonstrated 3 interventions that blocked the oscillation: (1) high-threshold calcium channel blockade (Figs. 1, 2), (2) fast sodium current blockade (Fig. 5) and (3) calcium-activated potassium current blockade (Fig. 9). The model helped to illustrate how all of these currents interacted to produce the oscillation (Fig. 11). During the oscillation, high fluxes through the high-threshold calcium channel (I_L_) and the calcium-activated potassium channel (I_AHP_) alternated to dominate the burst and interburst periods respectively. The dependence of the oscillation on the fast sodium current was a secondary effect, due to the preferential influx of calcium through high-threshold calcium channels during sodium spikes - a consequence of the rapid activation kinetics and high gain of I_L_ in the spike voltage range. Taken together, these results showed that the high-threshold calcium (I_L_) and calcium-activated potassium (I_AHP_) currents were critical for generating the intrinsic oscillation observed in local thalamic interneurons of the rat LGN.

**Figure 11:**
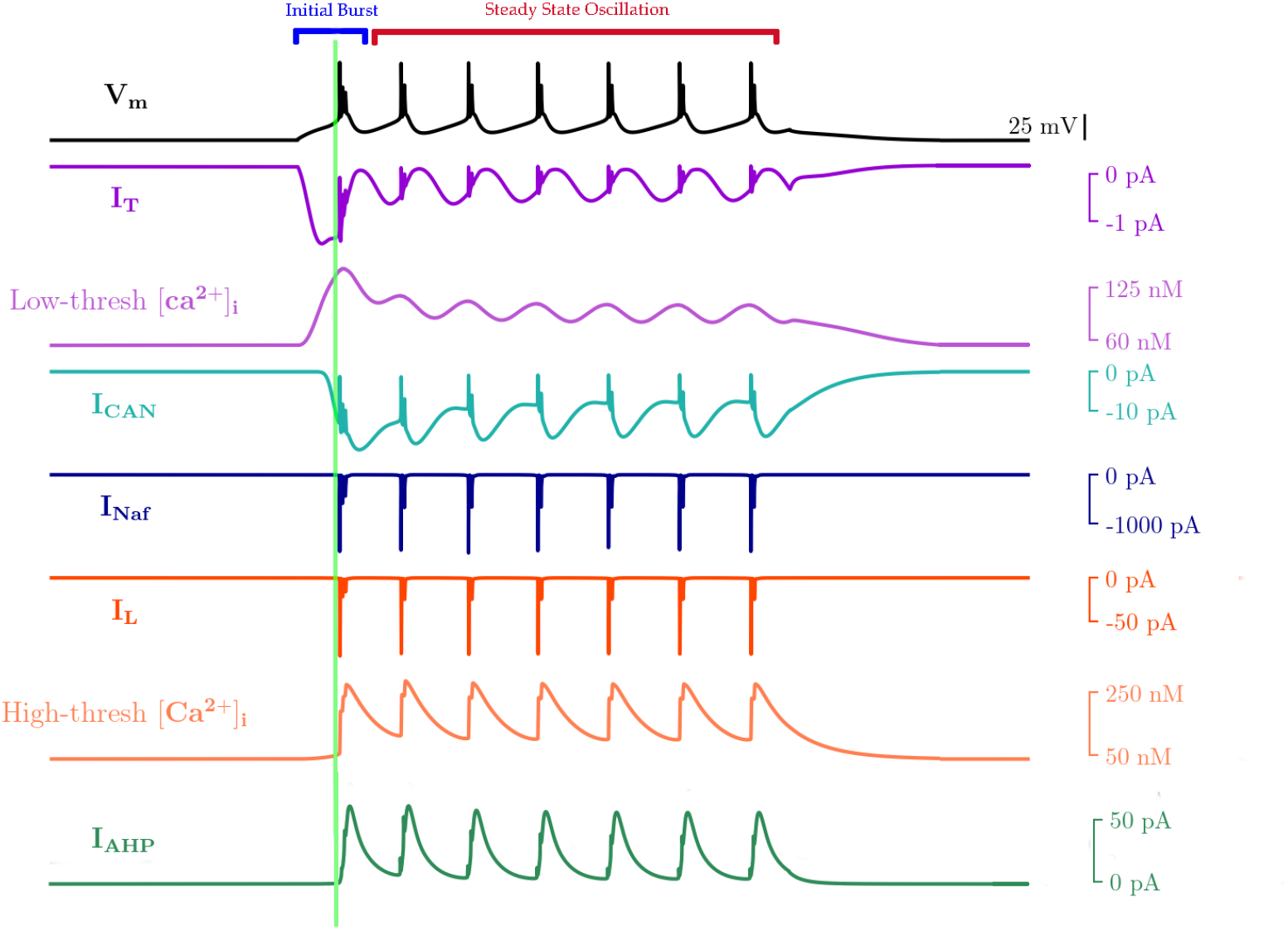
Origin of steady-state oscillatory bursting with 0.11 nA current clamp. Initial depolarization activates *I*_CAN_, *I*_T_, and feeds *I*_T_ [Ca^++^]_*i*_ pool, leading to initial burst (green line), terminated through hyperpolarization due to *I*_AHP_. Then, the continuing depolarization inactivates *I*_T_, leaving *I*_L_ to provide the depolarizing *I*_Ca_ during the repeated bursts. (V_m_: soma voltage; Currents: *I*_T_, *I*_L_: T-, L-type Ca^2+^ currents; *I*_CAN_: nonselective cation; *I*_Naf_: fast Na^+^; *I*_AHP_: Ca^2+^-activated K^+^; Ca^2+^ concentrations: separate pools served by *I*_T_ vs. *I*_L_)

Additionally, in our simulations, we noted that reductions in I_T_ density often resulted in delayed initial bursts (Fig. 3). Looking at how the currents interact in the model, we can see that I_T_ is the first conductance active prior to the initial burst, with I_CAN_ being activated second, presumably due to the calcium influx from I_T_ (Fig. 11). Recall also that *in silico* elimination of I_CAN_ did not abolish the oscillation, but instead also increased the time to first burst (Fig. 4). Given what we see from the model, we concluded that I_CAN_ provided a depolarizing boost, allowing the cell to produce the initial burst earlier than it would have with I_T_ alone. By using the model to examine the anatomy of the initial burst, we thus predict that initial depolarization from the low-threshold I_T_ current, with a depolarizing boost from I_CAN_, eventually leads the cell to fire a sodium spike, triggering a burst from I_L_ which then initiates the bursting oscillation (Fig. 11).

## 4 Discussion

Our use of detailed pharmacological blockade evidence represents an advance over existing models of this cell type (Bloomfield and Sherman, 1989; Perreault and Raastad, 2006; Briska et al., 2003). This approach helped us reduce concerns about degeneracy, a problem commonly encountered in modeling, where several different parameter sets can produce the same or similar outcomes (Taylor et al., 2009; Jelescu et al., 2016; Halnes et al., 2011; Huguenard and McCormick, 1992; Lederman et al., 2022). In certain contexts, parameter degeneracy can be considered a feature, since it can reflect a real variety found in nature (Neymotin et al., 2016; Edelman and Gally, 2001; Goaillard and Marder, 2021; Medlock et al., 2022). However, in this case, previous studies of thalamic interneurons indicated uncertainty regarding the role of voltage-gated calcium conductances and hyperpolarizing conductances, including I_AHP_ (Halnes et al., 2011; Zhu et al., 1999a) and I_A_ (Halnes et al., 2011; Zhu et al., 1999b), in sculpting single or repetitive bursting patterns. The tandem approach used here thus helped to resolve some of this uncertainty by pharmacologically identifying specific channel types that were crucial to the oscillation. Indeed, a different multicompartment model of this cell type manipulated the density of low-voltage-activated calcium channels (I_*T*_ ) and I_*AHP*_ to achieve periodic bursting in response to depolarizing current input (Halnes et al., 2011). Although this parameterization successfully evoked repetitive bursts, the resulting prediction regarding the importance of I_*T*_ in driving the intrinsic oscillation differed from the predictions generated by our pharmacological blockade experiments.

The behavior of low-vs. high-threshold calcium currents in thalamic interneurons differs from what is seen in thalamocortical relay cells and thalamic reticular cells. In the thalamic interneuron, the low-threshold calcium channels primarily contribute to the initial burst, but are not essential for periodic bursting - a phenomenon which instead utilizes the high-threshold L- and P-type calcium channels. This stands in contrast to the repetitive bursting mechanisms of both of the other cell types, which involve repeated contributions from low-threshold T-type calcium channels (Amarillo et al., 2015; Lee et al., 2014).

This type of prediction represents a distinct advantage of a paired *in vitro* and *in silico* approach. Electrophysiological experiments reduced the parameter space, providing valuable constraints to the modeling, which in turn was able to generate fine-grained analyses regarding the components and behavior of the system. Here, coupling physiology with modeling allowed us to first identify the most relevant conductances, then make detailed predictions regarding their interactions and contributions to different elements of the oscillation.

### 4.1 Limitations

1. We only briefly considered the role of dendritic channels, finding little change in somatic response with several variations. However, active dendritic conductances play an important role in determining the temporal and spatial properties of feedforward inhibition from local interneurons onto thalamocortical relay cells Cox and Beatty (2017). Synaptically-evoked calcium and sodium spikes in local thalamic interneurons can induce different modes of GABA release, which differ in their temporal and spatial properties (Acuna-Goycolea et al., 2008). Sodium spikes can induce fast, widespread elevations in intracellular calcium in both the axons and the dendrites, while calcium spikes preferentially spread through the dendrites, leading to slower, more sustained GABA release from the dendritic arbor alone (Acuna-Goycolea et al., 2008; Pressler and Regehr, 2013; Antal et al., 2010). These differences may have implications for the behavior of thalamic circuitry. Thus, active dendritic conductances may be of great importance in understanding the interplay of synaptic inputs, including those of the synaptic triad (Acuna-Goycolea et al., 2008; Cox and Beatty, 2017; Bickford, 2019; Guido, 2018).
2. As noted earlier (see Methods), experimental data collection was completed approximately 20 years prior to the development of our final simulation set. Thus, not all contemporary pharmacological manipulations were available, constraining the experimental techniques to those in use at the time of data collection.

### 4.2 Future Directions

Future phase plane and bifurcation analyses, comparable to that of Hindmarsh and Rose (Rose and Hindmarsh, 1989c,9,9), could provide further insight into the genesis of thalamic interneuron dynamics. Beyond determining which currents are necessary for the oscillation, these types of analyses would help establish if and how other currents work to perform additional functions, such as stabilizing the oscillation or modulating its range (Amarillo et al., 2015).

Due to ongoing activity *in vivo*, electrophysiology and drug responses will likely differ from those recorded *in vitro*(Holt et al., 1996; Fernandez et al., 2018; Belle et al., 2018). As recording technology develops further, *in vivo* studies of thalamic interneurons would be immensely valuable, as would the study of activation patterns in thalamic interneuron dendrites (Gao et al., 2021; Briska et al., 2003). Investigations in different animal models would also be important, given the large differences in the density of thalamic interneurons across species (Jager et al., 2021), with local thalamic interneurons being largely restricted to the LGN and comprising only *∼*6% of thalamic neurons in rodents, while making up *∼*30% in primates and having a more widespread distribution throughout thalamus (Jager et al., 2021; Braak and Bachmann, 1985; Arcelli et al., 1997).

Placing this cell into a network context would allow for a more complete understanding of thalamic and thalamocortical network dynamics (Bhattacharya et al., 2016; Lorincz et al., 2009; Dura-Bernal et al., 2023). In many brain areas, interneuron activity has been established as an important part of network rhythmogenesis Skinner and Saraga (2010). As with thalamocortical cells, a shift between tonic and burst firing modes in the thalamic interneuron could lead to differences in network oscillation patterns (Lorincz et al., 2009) via firing mode shifts (Fig. 10). Additionally, a major role of thalamic interneurons is the mediation of inputs to thalamus via the synaptic triads on distal dendrites (Bickford, 2019). Since this project was focused on uncovering the mechanism of the intrinsic oscillation, our current modeling does not address the dynamics occurring at these locations. However, the conclusions drawn here regarding the importance of high-threshold calcium conductances (I_L_) and the hyperpolarizing I_AHP_ current in generating the intrinsic oscillation may provide new targets for manipulating ascending thalamocortical network activity. These results may provide new targets for manipulating ascending thalamocortical network activity and help advance our understanding of both pathological (e.g. seizure activity (Paz and Huguenard, 2015)) and non-pathological activity in thalamocortical networks.

## Acknowledgments

The authors would like to thank Michael Hines and Ted Carnevale (Yale) for NEURON simulator support, Larry Eberle (SUNY Downstate) for Neurosim lab support, Tom Morse and Robert McDougal (Yale) for ModelDB support. Research supported by NIH R01DC012947, R01DC019979, P50MH109429, R01MH134118, R01NS128924, U24EB028998, ARL W911NF-22-2-0139, and NYS DOH SCIRB C38328GG.

## References

Acuna-Goycolea, C., Brenowitz, S. D., and Regehr, W. G. (2008). Active dendritic conductances dynamically regulate GABA release from thalamic interneurons. Neuron, 57(3):420–431.

Adams, D. J. and Berecki, G. (2013). Mechanisms of conotoxin inhibition of n-type (cav2.2) calcium channels. Biochimica et Biophysica Acta (BBA) - Biomembranes, 1828(7):1619–1628.

Allken, V., Chepkoech, J.-L., Einevoll, G. T., and Halnes, G. (2014). The subcellular distribution of t-type ca2+ channels in interneurons of the lateral geniculate nucleus. PLoS One, 9(9):e107780.

Amarillo, Y., Mato, G., and Nadal, M. S. (2015). Analysis of the role of the low threshold currents IT and ih in intrinsic delta oscillations of thalamocortical neurons. Front. Comput. Neurosci., 9:52.

Antal, M., Acuna-Goycolea, C., Pressler, R. T., Blitz, D. M., and Regehr, W. G. (2010). Cholinergic activation of M2 receptors leads to context-dependent modulation of feedforward inhibition in the visual thalamus. PLoS Biol., 8(4):e1000348.

Arcelli, P., Frassoni, C., Regondi, M. C., De Biasi, S., and Spreafico, R. (1997). GABAergic neurons in mammalian thalamus: a marker of thalamic complexity? Brain Res. Bull., 42(1):27–37.

Augustinaite, S., Yanagawa, Y., and Heggelund, P. (2011). Cortical feedback regulation of input to visual cortex: role of intrageniculate interneurons. J. Physiol., 589(Pt 12):2963–2977.

Avoli, M. (2012). A brief history on the oscillating roles of thalamus and cortex in absence seizures. Epilepsia, 53(5):779–789.

Babadi, B., Casti, A., Xiao, Y., Kaplan, E., and Paninski, L. (2010). A generalized linear model of the impact of direct and indirect inputs to the lateral geniculate nucleus. J. Vis., 10(10):22.

Banks, M. I., Pearce, R. A., and Smith, P. H. (1993). Hyperpolarization-activated cation current (ih) in neurons of the medial nucleus of the trapezoid body: voltage-clamp analysis and enhancement by norepinephrine and cAMP suggest a modulatory mechanism in the auditory brain stem. J. Neurophysiol., 70(4):1420–1432.

Bastos, A. M., Briggs, F., Alitto, H. J., Mangun, G. R., and Usrey, W. M. (2014). Simultaneous recordings from the primary visual cortex and lateral geniculate nucleus reveal rhythmic interactions and a cortical source for ?-band oscillations. J. Neurosci., 34(22):7639–7644.

Belle, A. M., Enright, H. A., Sales, A. P., Kulp, K., Osburn, J., Kuhn, E. A., Fischer, N. O., and Wheeler, E. K. (2018). Evaluation of in vitro neuronal platforms as surrogates for in vivo whole brain systems. Sci. Rep., 8(1):10820.

Bhattacharya, B. S., Bond, T. P., O’Hare, L., Turner, D., and Durrant, S. J. (2016). Causal role of thalamic interneurons in brain state transitions: A study using a neural mass model implementing synaptic kinetics. Front. Comput. Neurosci., 10:115.

Bhattacharya, B. S., Coyle, D., and Maguire, L. P. (2011). Alpha and theta rhythm abnormality in alzheimer’s disease: a study using a computational model. Adv. Exp. Med. Biol., 718:57–73.

Bickford, M. E. (2019). Synaptic organization of the dorsal lateral geniculate nucleus. Eur. J. Neurosci., 49(7):938–947.

Bloomfield, S. A. and Sherman, S. M. (1989). Dendritic current flow in relay cells and interneurons of the cat’s lateral geniculate nucleus. Proc. Natl. Acad. Sci. U. S. A., 86(10):3911–3914.

Bond, C. T., Herson, P. S., Strassmaier, T., Hammond, R., Stackman, R., Maylie, J., and Adelman, J. P. (2004). Small conductance ca2+-activated k+ channel knock-out mice reveal the identity of calcium-dependent afterhyperpolarization currents. J. Neurosci., 24(23):5301–5306.

Bonin, R. P., Zurek, A. A., Yu, J., Bayliss, D. A., and Orser, B. A. (2013). Hyperpolarization-activated current (in) is reduced in hippocampal neurons from gabra5-/-mice. PLoS One, 8(3):e58679.

Bonjean, M., Baker, T., Lemieux, M., Timofeev, I., Sejnowski, T., and Bazhenov, M. (2011). Corticothalamic feedback controls sleep spindle duration in vivo. J. Neurosci., 31(25):9124–9134.

Borg-Graham, L. (1991). Modelling the non-linear conductances of excitable membranes. In Wheal, H and Chad,, editor, Cellular Neurobiology: A Practical Approach, pages 247–275. Oxford University Press.

Braak, H. and Bachmann, A. (1985). The percentage of projection neurons and interneurons in the human lateral geniculate nucleus. Hum. Neurobiol., 4(2):91–95.

Briska, A. M., Uhlrich, D. J., and Lytton, W. W. (2003). Computer model of passive signal integration based on whole-cell in vitro studies of rat lateral geniculate nucleus. Eur. J. Neurosci., 17(8):1531–1541.

Budde, T., Munsch, T., and Pape, H. C. (1998). Distribution of l-type calcium channels in rat thalamic neurones. Eur. J. Neurosci., 10(2):586–597.

Chausson, P., Leresche, N., and Lambert, R. C. (2013). Dynamics of intrinsic dendritic calcium signaling during tonic firing of thalamic reticular neurons. PLoS One, 8(8):e72275.

Chen, Z., Wimmer, R. D., Wilson, M. A., and Halassa, M. M. (2015). Thalamic circuit mechanisms link sensory processing in sleep and attention. Front. Neural Circuits, 9:83.

Chow, R. H. (1991). Cadmium block of squid calcium currents. macroscopic data and a kinetic model. J. Gen. Physiol., 98(4):751–770.

Christina I. Schroeder, R. J. L. (2006). ?-Conotoxins GVIA, MVIIA and CVID: SAR and clinical potential. Mar. Drugs, 4(3):193.

Coulon, P., Budde, T., and Pape, H.-C. (2012). The sleep relay–the role of the thalamus in central and decentral sleep regulation. Pflugers Arch., 463(1):53–71.

Coulon, P., Herr, D., Kanyshkova, T., Meuth, P., Budde, T., and Pape, H.-C. (2009). Burst discharges in neurons of the thalamic reticular nucleus are shaped by calcium-induced calcium release. Cell Calcium, 46(5-6):333–346.

Cox, C. L. and Beatty, J. A. (2017). The multifaceted role of inhibitory interneurons in the dorsal lateral geniculate nucleus. Vis. Neurosci., 34:E017.

Crandall, S. R., Govindaiah, G., and Cox, C. L. (2010). Low-threshold ca2+ current amplifies distal dendritic signaling in thalamic reticular neurons. J. Neurosci., 30(46):15419–15429.

Crunelli, V., Cope, D. W., and Hughes, S. W. (2006). Thalamic t-type ca2+ channels and NREM sleep. Cell Calcium, 40(2):175–190.

Crunelli, V., Haby, M., Jassik-Gerschenfeld, D., Leresche, N., and Pirchio, M. (1988). Cl- - and k+-dependent inhibitory postsynaptic potentials evoked by interneurones of the rat lateral geniculate nucleus. J. Physiol., 399:153–176.

David, F., Schmiedt, J. T., Taylor, H. L., Orban, G., Di Giovanni, G., Uebele, V. N., Renger, J. J., Lambert, R. C., Leresche, N., and Crunelli, V. (2013). Essential thalamic contribution to slow waves of natural sleep. J. Neurosci., 33(50):19599–19610.

Destexhe, A., Contreras, D., Sejnowski, T. J., and Steriade, M. (1994a). A model of spindle rhythmicity in the isolated thalamic reticular nucleus. J. Neurophysiol., 72(2):803–818.

Destexhe, A., Mainen, Z. F., and Sejnowski, T. J. (1994b). Synthesis of models for excitable membranes, synaptic transmission and neuromodulation using a common kinetic formalism. J. Comput. Neurosci., 1(3):195–230.

Destexhe, A. and Sejnowski, T. J. (2003). Interactions between membrane conductances underlying thalamocortical slow-wave oscillations. Physiol. Rev., 83(4):1401–1453.

Dilger, E. K., Shin, H.-S., and Guido, W. (2011). Requirements for synaptically evoked plateau potentials in relay cells of the dorsal lateral geniculate nucleus of the mouse. J. Physiol., 589(Pt 4):919–937.

Dubin, M. W. and Cleland, B. G. (1977). Organization of visual inputs to interneurons of lateral geniculate nucleus of the cat. J. Neurophysiol., 40(2):410–427.

Dura-Bernal, S., Griffith, E. Y., Barczak, A., O’Connell, M. N., McGinnis, T., Moreira, J. V., Schroeder, C. E., Lytton, W. W., Lakatos, P., and Neymotin, S. A. (2023). Data-driven multiscale model of macaque auditory thalamocortical circuits reproduces in vivo dynamics. Cell reports, 42(11).

Dura-Bernal, S., Suter, B. A., Gleeson, P., Cantarelli, M., Quintana, A., Rodriguez, F., Kedziora, D. J., Chadderdon, G. L., Kerr, C. C., Neymotin, S. A., McDougal, R. A., Hines, M., Shepherd, G. M., and Lytton, W. W. (2019). NetPyNE, a tool for data-driven multiscale modeling of brain circuits. Elife, 8.

Edelman, G. M. and Gally, J. A. (2001). Degeneracy and complexity in biological systems. Proc. Natl. Acad. Sci. U. S. A., 98(24):13763–13768.

Fernandez, F. R., Rahsepar, B., and White, J. A. (2018). Differences in the electrophysiological properties of mouse somatosensory layer 2/3 neurons in vivo and slice stem from intrinsic sources rather than a Network-Generated high conductance state. eNeuro, 5(2).

Fogerson, P. M. and Huguenard, J. R. (2016). Tapping the brakes: Cellular and synaptic mechanisms that regulate thalamic oscillations. Neuron, 92(4):687–704.

Foxe, J. J. and Snyder, A. C. (2011). The role of Alpha-Band brain oscillations as a sensory suppression mechanism during selective attention. Front. Psychol., 2:154.

Galligan, J. J., Tatsumi, H., Shen, K. Z., Surprenant, A., and North, R. A. (1990). Cation current activated by hyperpolarization (IH) in guinea pig enteric neurons. Am. J. Physiol., 259(6 Pt 1):G966–72.

Gao, P. P., Graham, J. W., Zhou, W.-L., Jang, J., Angulo, S., Dura-Bernal, S., Hines, M., Lytton, W. W., and Antic, S. D. (2021). Local glutamate-mediated dendritic plateau potentials change the state of the cortical pyramidal neuron. J. Neurophysiol., 125(1):23–42.

Garcia, M. L., Knaus, H. G., Munujos, P., Slaughter, R. S., and Kaczorowski, G. J. (1995). Charybdotoxin and its effects on potassium channels. Am. J. Physiol., 269(1 Pt 1):C1–10.

Gent, T. C., Bandarabadi, M., Herrera, C. G., and Adamantidis, A. R. (2018). Thalamic dual control of sleep and wakefulness. Nat. Neurosci., 21(7):974–984.

Goaillard, J.-M. and Marder, E. (2021). Ion channel degeneracy, variability, and covariation in neuron and circuit resilience. Annu. Rev. Neurosci., 44:335–357.

Goh, J. W., Kelly, M. E., Pennefather, P. S., Chicchi, G. G., Cascieri, M. A., Garcia, M. L., and Kaczorowski, G. J. (1992). Effect of charybdotoxin and leiurotoxin I on potassium currents in bullfrog sympathetic ganglion and hippocampal neurons. Brain Res., 591(1):165–170.

Grimbert, F. and Faugeras, O. (2006). Bifurcation analysis of jansen’s neural mass model. Neural Comput., 18(12):3052–3068.

Guido, W. (2018). Development, form, and function of the mouse visual thalamus. J. Neurophysiol., 120(1):211–225.

Gurney, A. M., Tsien, R. Y., and Lester, H. A. (1987). Activation of a potassium current by rapid photochemically generated step increases of intracellular calcium in rat sympathetic neurons. Proc. Natl. Acad. Sci. U. S. A., 84(10):3496–3500.

Halnes, G., Augustinaite, S., Heggelund, P., Einevoll, G. T., and Migliore, M. (2011). A multi-compartment model for interneurons in the dorsal lateral geniculate nucleus. PLoS Comput. Biol., 7(9):e1002160.

Hines, M. (1993). NEURON — a program for simulation of nerve equations. In Eeckman, F. H., editor, Neural Systems: Analysis and Modeling, pages 127–136. Springer US, Boston, MA.

Hirsch, J. A., Wang, X., Sommer, F. T., and Martinez, L. M. (2015). How inhibitory circuits in the thalamus serve vision. Annu. Rev. Neurosci., 38:309–329.

Hodkinson, D. J., Wilcox, S. L., Veggeberg, R., Noseda, R., Burstein, R., Borsook, D., and Becerra, L. (2016). Increased amplitude of thalamocortical Low-Frequency oscillations in patients with migraine. J. Neurosci., 36(30):8026–8036.

Holt, G. R., Softky, W. R., Koch, C., and Douglas, R. J. (1996). Comparison of discharge variability in vitro and in vivo in cat visual cortex neurons. J. Neurophysiol., 75(5):1806–1814.

Horikawa, K. and Armstrong, W. E. (1988). A versatile means of intracellular labeling: injection of biocytin and its detection with avidin conjugates. J. Neurosci. Methods, 25(1):1–11.

Huguenard, J. R. and McCormick, D. A. (1992). Simulation of the currents involved in rhythmic oscillations in thalamic relay neurons. J. Neurophysiol., 68(4):1373–1383.

Jager, P., Moore, G., Calpin, P., Durmishi, X., Kita, Y., Salgarella, I., Wang, Y., Schultz, S. R., Brickley, S., Shimogori, T., and Delogu, A. (2021). Dual midbrain and forebrain origins of thalamic inhibitory interneurons. eLife, page e59272.

Jelescu, I. O., Veraart, J., Fieremans, E., and Novikov, D. S. (2016). Degeneracy in model parameter estimation for multi-compartmental diffusion in neuronal tissue. NMR Biomed., 29(1):33–47.

Kang, H.-W., Park, J.-Y., Jeong, S.-W., Kim, J.-A., Moon, H.-J., Perez-Reyes, E., and Lee, J.-H. (2006). A molecular determinant of nickel inhibition in cav3.2 t-type calcium channels. J. Biol. Chem., 281(8):4823–4830.

Kay, A. R. and Wong, R. K. (1987). Calcium current activation kinetics in isolated pyramidal neurones of the ca1 region of the mature guinea-pig hippocampus. J. Physiol., 392:603–616.

Kerschensteiner, D. and Guido, W. (2017). Organization of the dorsal lateral geniculate nucleus in the mouse. Vis. Neurosci., 34:E008.

Ketz, N. A., Jensen, O., and O’Reilly, R. C. (2015). Thalamic pathways underlying prefrontal cortex-medial temporal lobe oscillatory interactions. Trends Neurosci., 38(1):3–12.

Kim, H. R., Hong, S. Z., and Fiorillo, C. D. (2015). T-type calcium channels cause bursts of spikes in motor but not sensory thalamic neurons during mimicry of natural patterns of synaptic input. Front. Cell. Neurosci., 9:428.

Kitayama, M., Miyata, H., Yano, M., Saito, N., Matsuda, Y., Yamauchi, T., and Kogure, S. (2003). Ih blockers have a potential of antiepileptic effects. Epilepsia, 44(1):20–24.

Lakatos, P., O’Connell, M. N., Barczak, A., McGinnis, T., Neymotin, S., Schroeder, C. E., Smiley, J. F., and Javitt, D. C. (2020). The thalamocortical circuit of auditory mismatch negativity. Biological psychiatry, 87(8):770–780.

Lambert, N. A. and Wilson, W. A. (1996). High-threshold ca2+ currents in rat hippocampal interneurones and their selective inhibition by activation of GABA(B) receptors. J. Physiol., 492 (Pt 1):115–127.

Lansman, J. B., Hess, P., and Tsien, R. W. (1986). Blockade of current through single calcium channels by cd2+, mg2+, and ca2+. voltage and concentration dependence of calcium entry into the pore. J. Gen. Physiol., 88(3):321–347.

Lederman, D., Patel, R., Itani, O., and Rotstein, H. G. (2022). Parameter estimation in the age of degeneracy and unidentifiability. Sci. China Ser. A Math., 10(2):170.

Lee, J. H., Gomora, J. C., Cribbs, L. L., and Perez-Reyes, E. (1999). Nickel block of three cloned t-type calcium channels: low concentrations selectively block alpha1h. Biophys. J., 77(6):3034–3042.

Lee, S. E., Lee, J., Latchoumane, C., Lee, B., Oh, S.-J., Saud, Z. A., Park, C., Sun, N., Cheong, E., Chen, C.-C., Choi, E.-J., Lee, C. J., and Shin, H.-S. (2014). Rebound burst firing in the reticular thalamus is not essential for pharmacological absence seizures in mice. Proc. Natl. Acad. Sci. U. S. A., 111(32):11828–11833.

Lörincz, M. L., Crunelli, V., and Hughes, S. W. (2008). Cellular dynamics of cholinergically induced alpha (8-13 hz) rhythms in sensory thalamic nuclei in vitro. J. Neurosci., 28(3):660–671.

Lorincz, M. L., Kékesi, K. A., Juhász, G., Crunelli, V., and Hughes, S. W. (2009). Temporal framing of thalamic relay-mode firing by phasic inhibition during the alpha rhythm. Neuron, 63(5):683–696.

Lytton, W. W. and Sejnowski, T. J. (1991). Simulations of cortical pyramidal neurons synchronized by inhibitory interneurons. J. Neurophysiol., 66(3):1059–1079.

Marder, E. and Abbott, L. F. (1995). Theory in motion. Curr. Opin. Neurobiol., 5(6):832–840.

Matthews, R. T. and Lee, W. L. (1991). A comparison of extracellular and intracellular recordings from medial septum/diagonal band neurons in vitro. Neuroscience, 42(2):451–462.

McCormick, D. A. and Huguenard, J. R. (1992). A model of the electrophysiological properties of thalamocortical relay neurons. J. Neurophysiol., 68(4):1384–1400.

McCormick, D. A. and Pape, H. C. (1988). Acetylcholine inhibits identified interneurons in the cat lateral geniculate nucleus. Nature, 334(6179):246–248.

McDonough, S. I., Boland, L. M., Mintz, I. M., and Bean, B. P. (2002). Interactions among toxins that inhibit n-type and p-type calcium channels. J. Gen. Physiol., 119(4):313–328.

McDonough, S. I., Swartz, K. J., Mintz, I. M., Boland, L. M., and Bean, B. P. (1996). Inhibition of calcium channels in rat central and peripheral neurons by omega-conotoxin MVIIC. J. Neurosci., 16(8):2612–2623.

McFarlane, M. B. and Gilly, W. F. (1998). State-dependent nickel block of a high-voltage-activated neuronal calcium channel. J. Neurophysiol., 80(4):1678–1685.

Medlock, L., Sekiguchi, K., Hong, S., Dura-Bernal, S., Lytton, W. W., and Prescott, S. A. (2022). Multiscale computer model of the spinal dorsal horn reveals changes in network processing associated with chronic pain. J. Neurosci., 42(15):3133–3149.

Mrejeru, A., Wei, A., and Ramirez, J. M. (2011). Calcium-activated non-selective cation currents are involved in generation of tonic and bursting activity in dopamine neurons of the substantia nigra pars compacta. J. Physiol., 589(Pt 10):2497–2514.

Munsch, T., Budde, T., and Pape, H. C. (1997). Voltage-activated intracellular calcium transients in thalamic relay cells and interneurons. Neuroreport, 8(11):2411–2418.

Neymotin, S. A., McDougal, R. A., Bulanova, A. S., Zeki, M., Lakatos, P., Terman, D., Hines, M. L., and Lytton, W. W. (2016). Calcium regulation of HCN channels supports persistent activity in a multiscale model of neocortex. Neuroscience, 316:344–366.

Nielsen, K. J., Schroeder, T., and Lewis, R. (2000). Structure-activity relationships of omega-conotoxins at n-type voltage-sensitive calcium channels. J. Mol. Recognit., 13(2):55–70.

Nikonenko, I., Bancila, M., Bloc, A., Muller, D., and Bijlenga, P. (2005). Inhibition of t-type calcium channels protects neurons from delayed ischemia-induced damage. Mol. Pharmacol., 68(1):84–89.

Nimmrich, V. and Gross, G. (2012). P/Q-type calcium channel modulators. Br. J. Pharmacol., 167(4):741–759.

Oh, M. M., Power, J. M., Thompson, L. T., and Disterhoft, J. F. (2000). Apamin increases excitability of CA1 hippocampal pyramidal neurons. Neurosci. Res. Commun., 27(2):135–142.

Pape, H. C., Budde, T., Mager, R., and Kisvárday, Z. F. (1994). Prevention of ca(2+)-mediated action potentials in GABAergic local circuit neurones of rat thalamus by a transient k+ current. J. Physiol., 478 Pt 3:403–422.

Pape, H. C. and McCormick, D. A. (1995). Electrophysiological and pharmacological properties of interneurons in the cat dorsal lateral geniculate nucleus. Neuroscience, 68(4):1105–1125.

Parajuli, L. K., Fukazawa, Y., Watanabe, M., and Shigemoto, R. (2010). Subcellular distribution of α1G subunit of t-type calcium channel in the mouse dorsal lateral geniculate nucleus. J. Comp. Neurol., 518(21):4362–4374.

Pathak, D., Guan, D., and Foehring, R. C. (2016). Roles of specific kv channel types in repolarization of the action potential in genetically identified subclasses of pyramidal neurons in mouse neocortex. J. Neurophysiol., 115(5):2317–2329.

Paz, J. T. and Huguenard, J. R. (2015). Microcircuits and their interactions in epilepsy: is the focus out of focus? Nat. Neurosci., 18(3):351–359.

Perreault, M.-C. and Raastad, M. (2006). Contribution of morphology and membrane resistance to integration of fast synaptic signals in two thalamic cell types. J. Physiol., 577(Pt 1):205–220.

Porcaro, C., Di Lorenzo, G., Seri, S., Pierelli, F., Tecchio, F., and Coppola, G. (2017). Impaired brainstem and thalamic high-frequency oscillatory EEG activity in migraine between attacks. Cephalalgia, 37(10):915–926.

Pressler, R. T. and Regehr, W. G. (2013). Metabotropic glutamate receptors drive global persistent inhibition in the visual thalamus. J. Neurosci., 33(6):2494–2506.

Randall, A. and Tsien, R. W. (1995). Pharmacological dissection of multiple types of ca2+ channel currents in rat cerebellar granule neurons. J. Neurosci., 15(4):2995–3012.

Robinson, P. A., Rennie, C. J., Rowe, D. L., and O’Connor, S. C. (2004). Estimation of multiscale neurophysiologic parameters by electroencephalographic means. Hum. Brain Mapp., 23(1):53–72.

Rose, R. M. and Hindmarsh, J. L. (1989a). The assembly of ionic currents in a thalamic neurons: I The three-dimensional model. prslb, 237:267–288.

Rose, R. M. and Hindmarsh, J. L. (1989b). The assembly of ionic currents in a thalamic neurons: II The stability and state diagrams. prslb, 237:289–312.

Rose, R. M. and Hindmarsh, J. L. (1989c). The assembly of ionic currents in a thalamic neurons: III The seven-dimensional model. prslb, 237:313–334.

Ryu, P. D. and Randic, M. (1990). Low- and high-voltage-activated calcium currents in rat spinal dorsal horn neurons. J. Neurophysiol., 63(2):273–285.

Saalmann, Y. B. and Kastner, S. (2011). Cognitive and perceptual functions of the visual thalamus. Neuron, 71(2):209–223.

Schwindt, P. C., Spain, W. J., Foehring, R. C., Stafstrom, C. E., Chubb, M. C., and Crill, W. E. (1988). Multiple potassium conductances and their functions in neurons from cat sensorimotor cortex in vitro. J. Neurophysiol., 59(2):424–449.

Shen, J. B., Jiang, B., and Pappano, A. J. (2000). Comparison of l-type calcium channel blockade by nifedipine and/or cadmium in guinea pig ventricular myocytes. J. Pharmacol. Exp. Ther., 294(2):562–570.

Sitnikova, E., Hramov, A. E., Grubov, V. V., Ovchinnkov, A. A., and Koronovsky, A. A. (2012). On-off intermittency of thalamo-cortical oscillations in the electroencephalogram of rats with genetic predisposition to absence epilepsy. Brain Res., 1436:147–156.

Skinner, F. and Saraga, F. (2010). Single neuron models: Interneurons. In Cutsuridis, V., Graham, B., Cobb, S., and Vida, I., editors, Hippocampal Microcircuits: A Computational Modeler’s Resource Book, pages 399–422. Springer New York, New York, NY.

Svoboda, K. R. and Lupica, C. R. (1998). Opioid inhibition of hippocampal interneurons via modulation of potassium and hyperpolarization-activated cation (ih) currents. J. Neurosci., 18(18):7084–7098.

Swandulla, D. and Armstrong, C. M. (1989). Calcium channel block by cadmium in chicken sensory neurons. Proc. Natl. Acad. Sci. U. S. A., 86(5):1736–1740.

Taylor, A. L., Goaillard, J.-M., and Marder, E. (2009). How multiple conductances determine electrophysiological properties in a multicompartment model. J. Neurosci., 29(17):5573–5586.

Thoby-Brisson, M., Telgkamp, P., and Ramirez, J. M. (2000). The role of the hyperpolarization-activated current in modulating rhythmic activity in the isolated respiratory network of mice. J. Neurosci., 20(8):2994–3005.

Traub, R. D., Wong, R. K., Miles, R., and Michelson, H. (1991). A model of a CA3 hippocampal pyramidal neuron incorporating voltage-clamp data on intrinsic conductances. J. Neurophysiol., 66(2):635–650.

Vijayan, S. and Kopell, N. J. (2012). Thalamic model of awake alpha oscillations and implications for stimulus processing. Proc. Natl. Acad. Sci. U. S. A., 109(45):18553–18558.

Wang, X., Sommer, F. T., and Hirsch, J. A. (2011a). Inhibitory circuits for visual processing in thalamus. Curr. Opin. Neurobiol., 21(5):726–733.

Wang, X., Vaingankar, V., Soto Sanchez, C., Sommer, F. T., and Hirsch, J. A. (2011b). Thalamic interneurons and relay cells use complementary synaptic mechanisms for visual processing. Nat. Neurosci., 14(2):224–231.

Wang, X., Wei, Y., Vaingankar, V., Wang, Q., Koepsell, K., Sommer, F. T., and Hirsch, J. A. (2007). Feedforward excitation and inhibition evoke dual modes of firing in the cat’s visual thalamus during naturalistic viewing. Neuron, 55(3):465–478.

Wang, Y., Goodfellow, M., Taylor, P. N., and Baier, G. (2014). Dynamic mechanisms of neocortical focal seizure onset. PLoS Comput. Biol., 10(8):e1003787.

Wheeler, D. B., Randall, A., and Tsien, R. W. (1994). Roles of n-type and q-type ca2+ channels in supporting hippocampal synaptic transmission. Science, 264(5155):107–111.

Wisgirda, M. E. and Dryer, S. E. (1994). Functional dependence of ca(2+)-activated k+ current on L- and n-type ca2+ channels: differences between chicken sympathetic and parasympathetic neurons suggest different regulatory mechanisms. Proc. Natl. Acad. Sci. U. S. A., 91(7):2858–2862.

Zaman, T., Lee, K., Park, C., Paydar, A., Choi, J. H., Cheong, E., Lee, C. J., and Shin, H.-S. (2011). Cav2.3 channels are critical for oscillatory burst discharges in the reticular thalamus and absence epilepsy. Neuron, 70(1):95–108.

Zamponi, G. W., Bourinet, E., and Snutch, T. P. (1996). Nickel block of a family of neuronal calcium channels: subtype- and subunit-dependent action at multiple sites. J. Membr. Biol., 151(1):77–90.

Zhang, L. and McBain, C. J. (1995). Potassium conductances underlying repolarization and after-hyperpolarization in rat CA1 hippocampal interneurones. J. Physiol., 488 (Pt 3):661–672.

Zhang, Y., Mori, M., Burgess, D. L., and Noebels, J. L. (2002). Mutations in High-Voltage-Activated calcium channel genes stimulate Low-Voltage-Activated currents in mouse thalamic relay neurons. J. Neurosci., 22(15):6362–6371.

Zhu, J. and Heggelund, P. (2001). Muscarinic regulation of dendritic and axonal outputs of rat thalamic interneurons: a new cellular mechanism for uncoupling distal dendrites. J. Neurosci., 21(4):1148–1159.

Zhu, J. J. (1996). Functional role of local GABAegic interneurons in thalamic oscillations. PhD thesis, University of Wisconsin, Madison.

Zhu, J. J. and Lo, F. S. (1999). Three GABA receptor-mediated postsynaptic potentials in interneurons in the rat lateral geniculate nucleus. J. Neurosci., 19(14):5721–5730.

Zhu, J. J., Lytton, W. W., Xue, J. T., and Uhlrich, D. J. (1999a). An intrinsic oscillation in interneurons of the rat lateral geniculate nucleus. J. Neurophysiol., 81(2):702–711.

Zhu, J. J. and Uhlrich, D. J. (1997). Nicotinic receptor-mediated responses in relay cells and interneurons in the rat lateral geniculate nucleus. Neuroscience, 80(1):191–202.

Zhu, J. J. and Uhlrich, D. J. (1998). Cellular mechanisms underlying two muscarinic receptor-mediated depolarizing responses in relay cells of the rat lateral geniculate nucleus. Neuroscience, 87(4):767–781.

Zhu, J. J., Uhlrich, D. J., and Lytton, W. W. (1999b). Burst firing in identified rat geniculate interneurons. Neuroscience, 91(4):1445–1460.

Zhu, J. J., Uhlrich, D. J., and Lytton, W. W. (1999c). Properties of a hyperpolarization-activated cation current in interneurons in the rat lateral geniculate nucleus. Neuroscience, 92(2):445–457.

